# Genome-wide sequence and expression analysis of the NAC transcription factor family in polyploid wheat

**DOI:** 10.1101/141747

**Authors:** Philippa Borrill, Sophie A. Harrington, Cristobal Uauy

**Affiliations:** Department of Crop Genetics, John Innes Centre, Norwich Research Park, NR4 7UH, UK

**Keywords:** wheat, transcription factors, NAC, phylogenetics, gene expression

## Abstract

Transcription factors are vital in plants to regulate gene expression in response to environmental stimuli and to control developmental processes. In this study, we annotated and classified transcription factors in polyploid bread wheat into gene families and explored the NAC family in detail. We combined phylogenetic analysis and transcriptome analysis, using publicly available RNA-seq data, to characterize the NAC gene family and provide hypotheses for putative functions of many NAC transcription factors. This study lays the groundwork for future studies on transcription factors in wheat which may be of great agronomic relevance.

**ABSTRACT:** Many important genes in agriculture correspond to transcription factors which regulate a wide range of pathways from flowering to responses to disease and abiotic stresses. In this study, we identified 5,776 transcription factors in hexaploid wheat (*Triticum aestivum*) and classified them into gene families. We further investigated the NAC family exploring the phylogeny, C-terminal domain conservation and expression profiles across 308 RNA-seq samples. Phylogenetic trees of NAC domains indicated that wheat NACs divided into eight groups similar to rice (*Oryza sativa*) and barley (*Hordeum vulgare*). C-terminal domain motifs were frequently conserved between wheat, rice and barley within phylogenetic groups, however this conservation was not maintained across phylogenetic groups. We explored gene expression patterns across a wide range of developmental stages, tissues, and abiotic stresses. We found that more phylogenetically related NACs shared more similar expression patterns compared to more distant NACs. However, within each phylogenetic group there were clades with diverse expression profiles. We carried out a co-expression analysis on all wheat genes and identified 37 modules of co-expressed genes of which 23 contained NACs. Using GO term enrichment we obtained putative functions for NACs within co-expressed modules including responses to heat and abiotic stress and responses to water: these NACs may represent targets for breeding or biotechnological applications. This study provides a framework and data for hypothesis generation for future studies on NAC transcription factors in wheat.

## INTRODUCTION

Transcription factors (TFs) by virtue of their role in activating or repressing gene expression, regulate many biological processes. They are particularly important to agriculture because TFs have been identified to be the causal genes underlying agronomic traits including flowering time, nutrient content, and stress responses (Yan et al. 2003; Uauy et al. 2006; Jensen and Skriver 2014). As such, identifying and characterizing the TFs in crops provides an important first step to engineer strategies for the improvement of agriculturally important traits.

Wheat is the most widely grown crop globally, providing roughly 20% of the daily calorific intake and 25% of protein intake worldwide (www.fao.org/faostat).The economic importance of wheat is also great, comprising over 40% of global cereal trade (FAO 2017). Twin pressures of increasing global population and changing climatic conditions make it ever more urgent that novel wheat varieties are developed which have improved yield potential, end-use quality, and increased tolerances to biotic and abiotic stresses, such as drought and heat.

Of the many TF families, the plant-specific NAC family has been shown to regulate several biological processes in wheat. Named after the first three such TFs identified (NAM, ATAF1/2 (Souer et al. 1996), and CUC2 (Aida et al. 1997)), the NAC TF family is characterized by a highly conserved NAC domain, typically at the N-terminal region, often followed by an intrinsically disordered transcriptional regulatory domain at the C-terminal region which is poorly conserved (Ernst et al. 2004; Olsen et al. 2005; Xie et al. 2000). The NAC domain is well characterized, and is required for protein-DNA interactions (Welner et al. 2012; Xie et al. 2000) and protein dimerization (Ernst et al. 2004). In wheat, NAC TFs are known to be involved in processes such as senescence and nutrient remobilization (Uauy et al. 2006; Zhao et al. 2015) as well as responses to abiotic and biotic stresses, ranging from stripe rust (Feng et al. 2014; Xia et al. 2010a; Xia et al. 2010b; Wang et al. 2015) to abiotic stresses including drought (Huang et al. 2015; Tang et al. 2012; Xue et al. 2006; Mao et al. 2014; Mao et al. 2012) and salt tolerance (Huang et al. 2015; Mao et al. 2014; Mao et al. 2012). The phylogenetic relationships of NAC TFs in different species have been identified and used to characterize evolutionarily-conserved groupings of NAC TFs (Ooka et al. 2003; Pereira-Santana et al. 2015; Shen et al. 2009). However, until recently such an analysis was hindered in wheat due to the lack of a high-quality reference genome sequence and a comprehensive set of gene models.

Recent advances in wheat genomics now provide the opportunity to characterize TF families much more completely in wheat (Uauy 2017). In this study, we used the recently published high quality TGAC gene models (Clavijo et al. 2017) to annotate all characterized TF families in wheat, and compare their abundance with other previously characterized crop species and wild relatives of wheat. We focused on the NAC TF family to understand the evolutionary relationships within the family itself and global expression patterns using large scale RNA-seq studies (Borrill et al. 2016; Clavijo et al. 2017) and co-expression networks. The analyses presented in this study allow novel hypotheses to be generated to predict TF function and pave the way for future functional characterization.

## MATERIALS AND METHODS

### Annotation of transcription factors

We downloaded the protein sequences for the gene models produced for the TGAC wheat assembly (Clavijo et al. 2017) from EnsemblPlants release-32 (Bolser et al. 2015) (http://plants.ensembl.org/index.html). These contained 249,547 transcripts corresponding to 195,864 genes of which 104,091 were high and 91,773 low confidence. We used these sequences to identify putative TFs using three methods for both high and low confidence genes.

#### 1. BLAST-based approach

We downloaded the protein sequences of TFs annotated in PlantTFDBv3.0 (Jin et al. 2014) fo*rAegilops tauschii, Hordeum vulgare, Oryza sativa* subsp. *japonica, Oryza sativa* subsp. *indica, Triticum urartu, Triticum aestivum* (from ESTs Unigene Build #63). We performed a blastp analysis of these protein sequences against the TGAC wheat protein sequences downloaded from EnsemblPlants with the parameter-max_target_seqs 10 to retrieve the top ten hits. We combined the BLAST results from each of the six species and removed duplicate genes.

#### 2. EnsembI orthologues based approach

We used EnsemblPlants Biomart to download wheat orthologues to the TFs in five species (*Aegilops tauschii, Hordeum vulgare, Oryza sativa* subsp. *japonica, Oryza sativa* subsp. *indica, Triticum urartu*) which were available on EnsemblPlants and annotated in PlantTFDBv3.0. For *Oryza sativa* subsp. *japonica* (before downloading wheat orthologues) we converted the MSU nomenclature rice gene identifiers from PlantTFDB to RAP rice gene identifiers which were compatible with EnsemblPlants using the RAPD converter http://rapdb.dna.affrc.go.ip/tools/converter/run. This step retained 1,816 RAP genes out of 1,859 MSU genes originally identified by PlantTFDB.

#### 3. TGAC functional annotation approach

We searched the functional annotation available for the TGAC wheat assembly (Clavijo et al. 2017) for all genes with PFAMs associated with TFs. The PFAMs associated with TFs were obtained from PlantTFDB.

### Generating a combined list of transcription factors

To generate a reliable list of TFs for wheat we combined the lists of genes identified by the blastp, EnsembI orthologue and functional annotation approaches. This included 9,416 genes (13,325 transcripts). This list may include genes which are not TFs in wheat due to changes to their sequences from their orthologues in the other monocot species or because genes with certain combinations of PFAM domains are known not to act as TFs (Jin et al. 2014). Therefore, we ran the 13,325 transcripts identified through the PlantTFDBv3.0 prediction server http://planttfdbv3.cbi.pku.edu.cn/prediction.php in batches of 1,000 genes. This resulted in the annotation of in total 7,415 genes (10,303 transcripts) of which 5,776 genes (8,609 transcripts) were from high confidence gene models. PlantTFDBv3.0 also assigned TFs to TF families.

### NAC transcription factor homoeologs and orthologs

From the list of TFs identified we extracted genes which were classified as NACs by PlantTFDBv3.0. For further analysis, we selected only NACs with high confidence gene models (453/574). For these 453 high confidence NACs genes, we downloaded information about wheat homoeologs from EnsemblPlants Biomart and grouped them into triads (A, B and D genome homoeologs). Rice (*Oryza sativa* subsp. *japonica*) and barley orthologs were identified by reciprocal BLAST of coding sequences. If the reciprocal BLAST did not identify the same pair of genes in both directions, they were not considered orthologs.

### Phylogenetic tree generation and NAC group assignment

We aligned the NAC protein sequences with Clustal Omega v1.2.0 (Sievers et al. 2011) using default settings. We kept only the NAC domain from the start of sub-domain A to the end of sub-domain E (Ooka et al. 2003; Uauy et al. 2006) to create phylogenetic trees for wheat, barley and rice NACs. After manual inspection, we found that a few regions within the NAC domain alignment were poorly conserved with amino acids only present in a few sequences. For this reason we only retained amino acid positions which were present in at least 10 % of sequences. We removed any sequences which did not contain any NAC domain sequence. We used RAxML v8.2.1 (Stamatakis 2014) to create maximum likelihood phylogenetic trees using the auto setting to detect the best protein model, 100 maximum likelihood searchers and 100 rapid bootstraps.

The barley and rice NACs had already been assigned to groups a-h in Christiansen et al. (2011) and Shen et al. (2009), respectively. Wheat genes which were phylogenetically grouped with barley or rice genes with a group classification were assigned to the appropriate group. In cases where the specific barley or rice orthologue belonged to a group dissimilar to the rest of the clade the wheat genes were not assigned to a group (23 genes). In total 430 wheat genes were assigned to a group. Figures with the groups alongside the NAC phylogeny were created using iTOL (Letunic and Bork 2016). We re-ran RAxML to make an individual phylogeny for groups a-h for wheat NACs and separately wheat, barley and rice NACs.

### C-terminal domain (CTD) motif discovery

We carried out *de novo* analysis of motifs in the a-h NAC TF groups using the MEME program (version 4.9.1)(Bailey et al. 2009). For each group, a maximum of 10 motifs were identified that occurred in all sequences and were between 5 and 20 residues long. From these motifs, we considered the most significant motif for further analysis, as well as additional significant motifs that shared sequence similarities with previously defined motifs (Ooka et al. 2003; Pereira-Santana et al. 2015; Shen et al. 2009).

To complement the *de novo* analysis, we screened all the wheat, barley, and rice NACs for motifs that were previously characterized (Ooka et al. 2003). A background amino acid frequency for wheat was obtained from the full set of peptide sequences from the TGAC gene models. We converted the motifs i-xiii from Ooka et al. (2003) into Regex expressions, and then converted into MEME motif format using IUPAC2MEME (v 4.9.1) from the MEME suite (Table S1). Using these motifs and the wheat amino acid background frequencies we searched all genes in the set with FIMO (v 4.9.1) from the MEME suite. In some cases, the Ooka groupings contained more than one motif (groups ii, iv, and ix; Table S1). Genes were considered part of group ii or group iv if at least one of the motifs was present. However, as the motifs from group ix were already split to form groups x and xi, only genes that contained both ix motifs were assigned to the ix group. Plots of the CTD motifs alongside the NAC phylogeny were created using iTOL (Letunic and Bork 2016).

### Gene expression analysis

We downloaded count and transcript per million (tpm) gene expression values for previously mapped RNA-seq samples from www.wheat-expression.com (Borrill et al. 2016; Clavijo et al. 2017). We excluded samples from cytogenetic stocks (e.g. nullitetrasomic lines) and from synthetic hexaploid wheat. This resulted in 308 RNA-seq samples from 15 individual studies being included in our analysis. We collated per transcript expression levels into per gene expression levels using the R package tximport v1.0.3 (Soneson et al. 2015). We filtered the data to only keep genes whose expression was over 0.5 tpm in at least three samples to eliminate very low expressed genes. We also filtered the data to exclude low confidence genes. We generated plots of phylogenetic trees with heatmaps of gene expression using the R package ggtree v1.4.20 (Yu et al. 2017).

### Co-expression analysis

We carried out co-expression analysis using the R package WGCNA v1.51 (Langfelder and Horvath 2008). We used the function pickSoftThreshold to calculate that a soft-threshold power of 6 was appropriate for a signed hybrid network for our 308 samples. Due to the large number of genes in our analysis (91,403) we used the blockwiseModules method to calculate the co-expression network in two blocks using the parameters maxPOutliers = 0.05, mergeCutHeight =0.15, deepSplit =2, minModuleSize = 30, networkType= “signed hybrid”, maxBlockSize = 46000, corType=“bicor”, corOptions = “use = ‘p’, maxPOutliers = 0.05”.

### Gene Ontology (GO) enrichment analysis

We used the R package GOseq v1.26.0 (Young et al. 2010) to determine whether GO terms were enriched within each co-expression module. We used Revigo (Supek et al. 2011) to summarize GO term enrichment for GO terms over-represented with a Benjamin Hochberg adjusted p-value <0.05.

### Data availability

The supplemental files contain the following data:

Table S1. NAC protein C-terminal domain motifs identified by Ooka et al. 2003.

Table S2. Wheat transcription factor family genes with gene model confidence levels.

Table S3. Wheat transcription factor distribution across chromosomes.

Table S4. Wheat, barley and rice NAC orthologues.

Table S5. C-terminal domain motifs per gene for wheat, barley and rice.

Table S6. *De novo* motif discovery in NAC groups.

Table S7. Gene and transcription factor module allocation by WGCNA co-expression analysis.

Table S8. Most over represented biological process GO terms in co-expression modules. Figure S1. Maximum likelihood phylogeny of wheat, barley and rice NAC transcription factor proteins constructed using the NAC domain.

Figure S2. Extended version of Figure 3, showing conserved C-terminal domains in wheat, rice and barley NAC transcription factors.

Figure S3. Extended version of Figure 4, showing gene expression of wheat NAC transcription factors in the context of the phylogeny.

Interactive trees for Figures 2, 3, S1 and S2 are available at http://itol.embl.de/shared/sophieharrington

## RESULTS

### Wheat transcription factors identified in the TGAC assembly

In total we annotated 5,776 high confidence genes as TFs in wheat which is a three-fold increase compared to the previous wheat TF annotation available from PlantTFDB (Table 1). We identified on average 5.1 times more TFs than in other diploid Triticeae species. However, for rice only 3.1 times more TFs were identified, as would be expected for a comparison between a diploid and hexaploid species. The incomplete nature of the Triticeae species’ genomes compared to the highly contiguous genome assemblies of rice may explain the higher than expected ratio to monocots other than rice. The annotation of low confidence genes was also carried out and a complete set of TFs in wheat is available in Table S2.

**Table 1.** Comparison of TFs identified in monocot species.

| Species | Ploidy | Transcription factor |  |  |
| --- | --- | --- | --- | --- |
|  |  | Transcripts<sup>1</sup> | Genes | Families |
| *Oryza sativa* subsp. *indica* | 2x | 1,891 | 1,891 | 56 |
| *Oryza sativa* subsp. *japonica* (MSU) | 2x | 2,408 | 1,859 | 56 |
| *Hordeum vulgare* | 2x | 2,621 | 1,198 | 56 |
| *Aegilops tauschii* | 2x | 1,439 | 1,439 | 55 |
| *Triticum urartu* | 2x | 888 | 888 | 50 |
| *Triticum aestivum* (ESTs Unigene #63) | 6x | 1,940 | 1,940 | 56 |
| *Triticum aestivum* (TGAC assembly high confidence genes) | 6x | 8,609 | 5,776 | 56 |

<sup>1</sup> Values are from PlantTFDB for all species, except *T. aestivum* TGAC assembly which is from this study.

We found that distribution of TF families was similar between wheat, barley and rice (Figure 1) with the largest families in all three species being bHLH and the smallest being STAT. In general, wheat had approximately three times more genes in each family than rice (Figure 1D, blue line). The only exceptions were the B3 and HB-other gene families which were enriched in wheat with five times as many genes as in rice (χ^2^ test p<0.001 and p=0.048 respectively). The FAR1 family was under-represented in wheat with only 2.5 times as many genes as in rice (χ^2^ test p=0.037). Compared to barley, most TF families had more members in wheat (Fig 1D, red line) which may be due to the incomplete nature of the barley genome.

**Figure 1.**
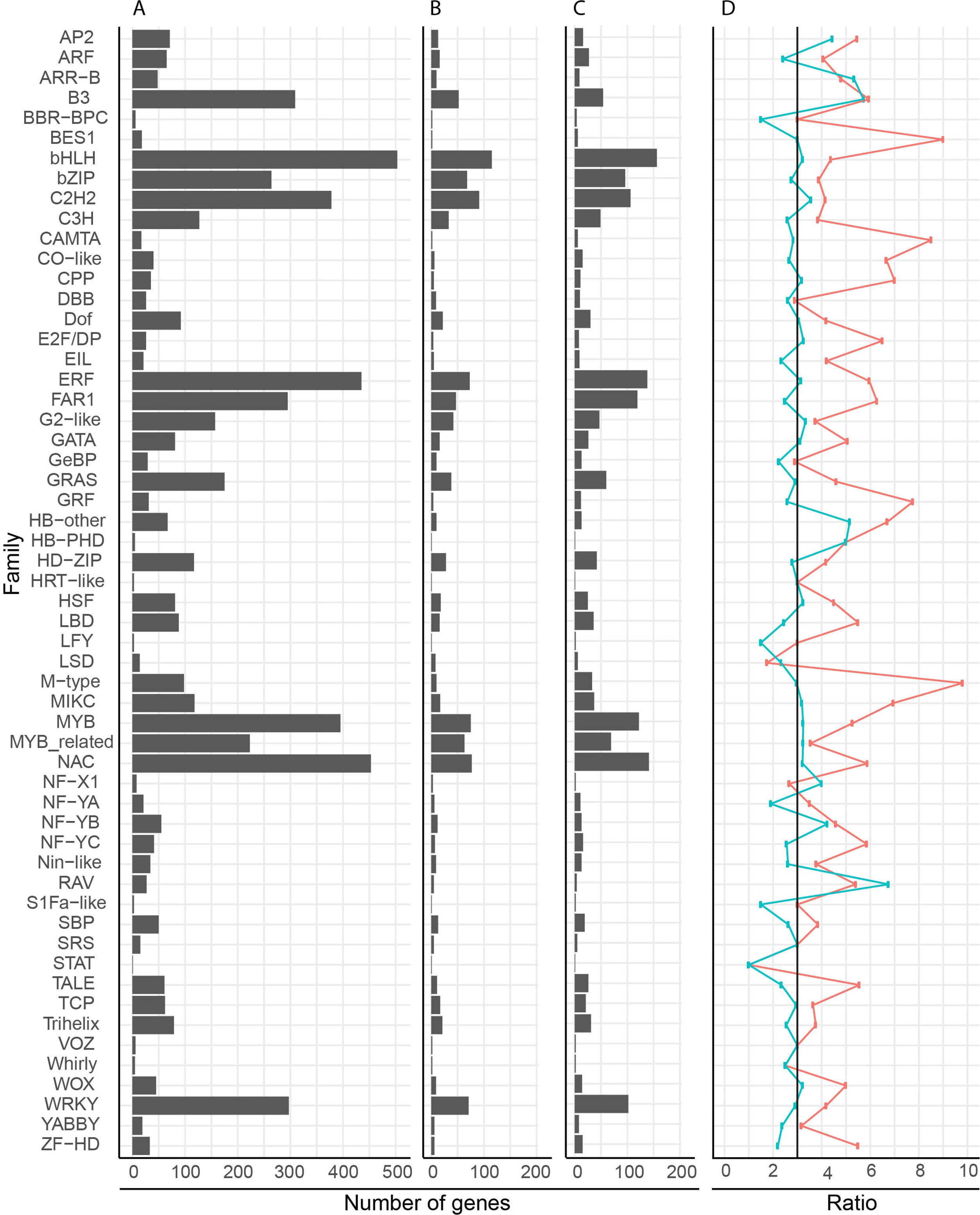
Comparison of genes identified per transcription factor family in wheat, barley and rice. The number of genes in each family for A) wheat, B) barley, C) rice. D) The ratio of wheat to barley (red) and wheat to rice (blue). In panel D) the expected ratio (3:1) is indicated by a black line. Barley and rice data were obtained from PlantTFDBv3.0.

We found that TFs were not distributed equally across all chromosomes with group 1 and group 6 having an average of 223 and 206 TFs per homoeolog, whereas group 3 and 5 had 300 and 304 TFs per homoeolog, respectively (Table S3). Individual TF families differed from the global averages, for example NAC TFs were most frequent on chromosome groups 2 and 7 and least frequent on groups 1 and 6, whereas WRKY TFs were most frequent on groups 1 and 3 and least frequent on groups 4 and 6.

### The NAC transcription factor family in wheat, barley and rice

We decided to focus our analysis on the NAC family of TFs which is known to be involved in a range of agronomically relevant processes including abiotic and biotic stress responses. In total we identified 453 NACs with high confidence gene models using the PlantTFDBv3.0 classifications. We grouped the NACs into homoeologous groups and identified their barley and rice orthologues by reciprocal blast (Table S4). To understand more about NAC evolution in wheat we generated a phylogenetic tree for wheat, barley and rice NACs using their NAC domains (Figure 2). We used the closest related barley and rice NACs to assign wheat NACs into eight main groups (a-h) as proposed by Shen et al. (2009) (see Figure S1).

**Figure 2.**
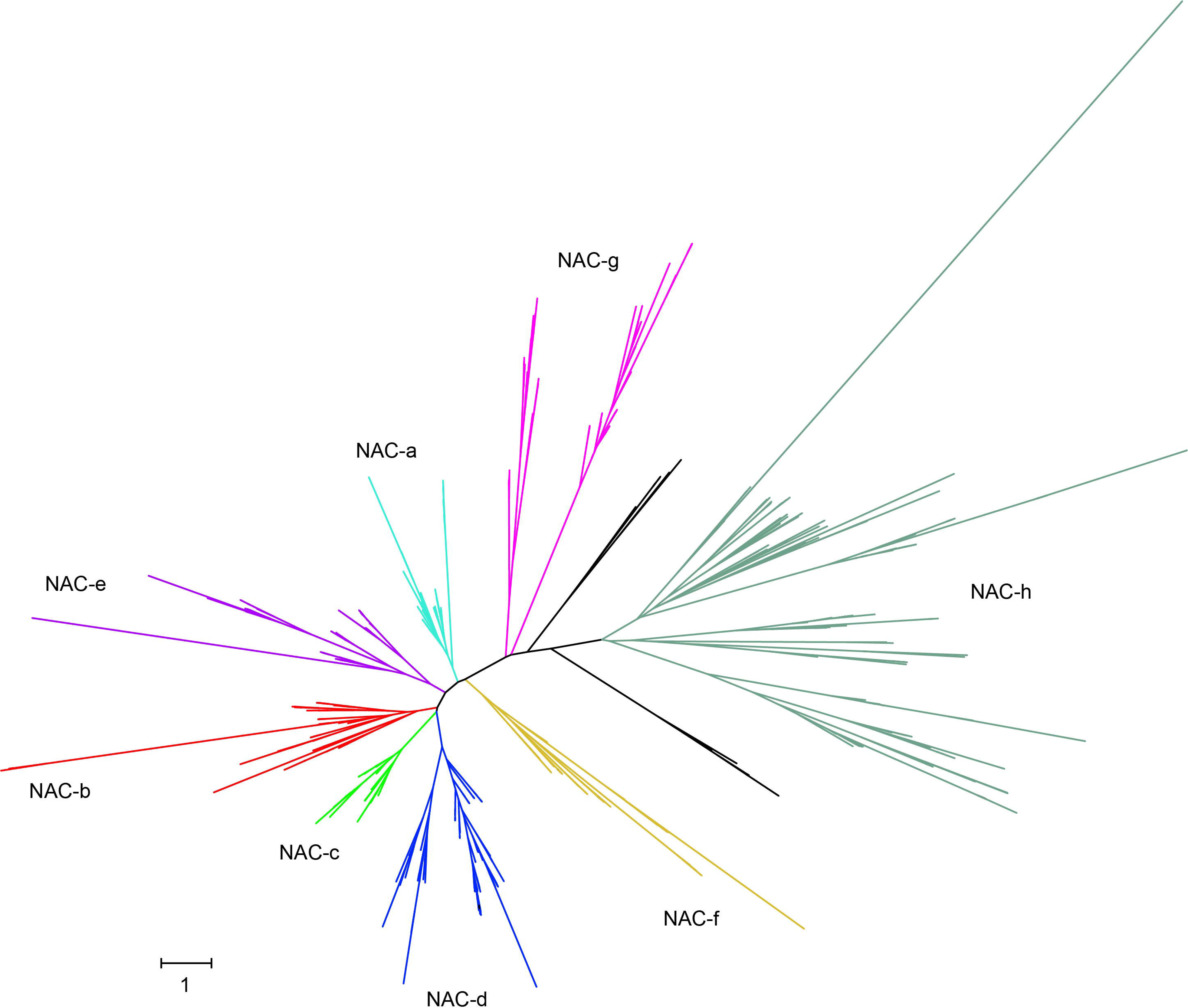
Maximum likelihood phylogeny of 667 NAC proteins from wheat, rice and barley. The phylogeny was constructed using only the NAC domain. NAC groups a-h were assigned according to rice and barley orthologues. In cases where the group assigned to a rice or barley gene conflicted with the overall tree topology no group was assigned (black branches). Details of individual genes are presented in Supplementary Table S4 and Figure SI.

In total 430 NACs were assigned to groups whilst 23 NACs could not be assigned to a group (either the NAC group was different for a particular protein compared to the rest of the clade or there was no clear rice or barley orthologue). As expected each group had in general three times more genes in wheat than in rice and barley. However, wheat has a reduced group f with only 13 genes compared to the 10 genes found in rice (χ^2^ p=0.001), but not compared to barley. Groups e, g and h are significantly enlarged in wheat compared to barley (χ^2^ p=0.04, p=0.04 and p<0.001 respectively) however the numbers of genes in each of these groups is lower in barley than in rice which suggests this trend is due to the incomplete barley genome rather than a true enrichment in wheat.

We also investigated the less well characterized C-terminal domain (CTD) which is proposed to be a transcriptional activator or repressor (Tran et al. 2004; Yamaguchi et al. 2010; Kim et al. 2007). We found that previously identified CTD motifs (Ooka et al. 2003) were generally conserved between homoeologs and were often conserved in specific clades within phylogenetic groups of wheat, barley and rice NACs (Figure 3, Figure S2, Table S5, http://itol.embl.de/shared/sophieharrington). We found that 10 out of the 13 motifs previously identified were present in wheat, rice and barley NACs. In general, each motif was predominantly found in one or two groups (e.g. motifs ii, v and vi were only in group a; motif vii in groups b and g; motif viii in group b). However, motif xiii was found in proteins belonging to all groups. The presence of motifs was not equally distributed between the groups with relatively few motifs in e, g and h and high frequency of motifs in c, d and f. *De novo* motif discovery identified significant motifs shared by all genes within each group (Table S6). Of these motifs, six had been previously identified as NAC CTD motifs (Ooka et al. 2003; Pereira-Santana et al. 2015; Shen et al. 2009), while three represent novel motifs.

**Figure 3.**
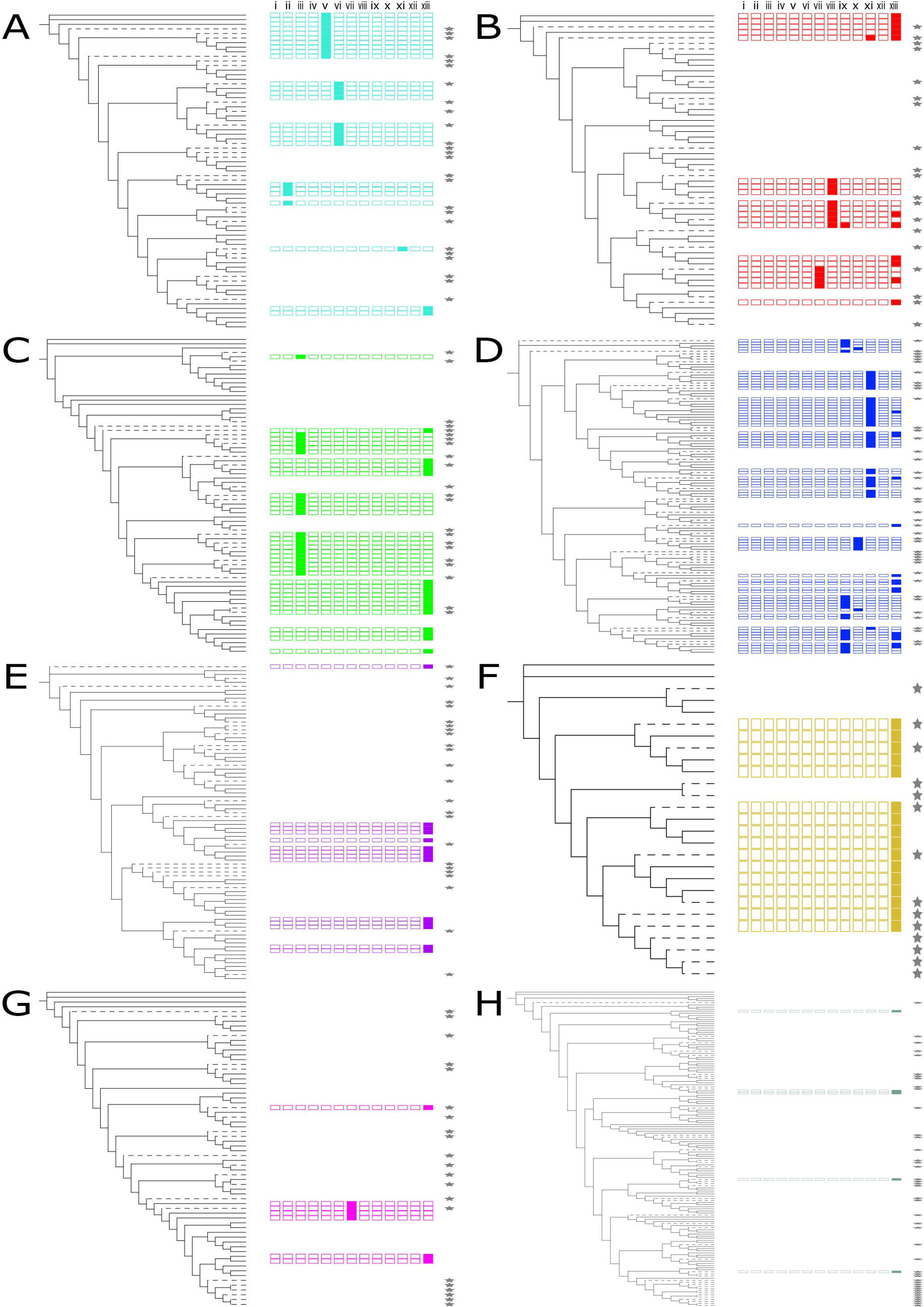
Conserved domains in the CTD of NAC transcription factors arranged by phylogenetic position. Known CTD motifs are shown alongside the wheat, barley, and rice NAC TF for each group a-h (A-H), colored in accordance with Figure 2. Branches corresponding to wheat NAC TFs are solid black; those for rice and barley NAC TFs are dashed. Motifs are shown as boxes, matching (left to right) motifs i – xiii from (Ooka et al. 2003). Motifs that are present in each protein (p-value < 0.05, q-value < 0.05) are shown by a solid-colored box, while absent motifs are shown by an empty outlined box. Genes with no significant motifs are shown with empty space. Barley and rice genes are indicated by the presence of a star to the right of the CTD motifs. Details are presented in Supplementary Table S5, and the full phylogenetic tree is presented in Figure S2.

### NAC expression patterns relate to phylogenetic position

To explore the expression patterns of NAC TFs we used publicly available gene expression data for 15 studies comprising 308 individual RNA-seq samples (Borrill et al. 2016; Clavijo et al. 2017). These samples included diverse developmental stages, tissues and stress conditions including both biotic and abiotic stresses. We filtered the NAC genes to retain only genes expressed at over 0.5 tpm in at least three samples. Within the phylogenetic groups a-h there were 430 NACs of which 356 passed this threshold. In most groups the vast majority of NAC genes were expressed, however in group h only 50 % of NACs were expressed in the conditions represented by the 308 RNA-seq samples.

We found that in general homoeologs shared similar expression patterns across samples (Figure 4, Figure S3). Gene expression patterns were more similar for genes found within the same phylogenetic group compared to genes in other groups. However, within each phylogenetic group gene expression patterns were more highly conserved within closely related clades than across the whole group. These conserved expression clades often showed expression specific to particular tissues or environmental conditions. For example in group d, 18 genes form a subclade which is predominantly expressed in the grain and the endosperm (Figure 4D, uppermost genes) and in group c, 20 genes form a clade which shows strong expression in spikelets which is not seen in other group c genes (Figure 4C, middle). We did not observe a correlation between expression patterns and the presence of specific CTDs (data not shown).

**Figure 4.**
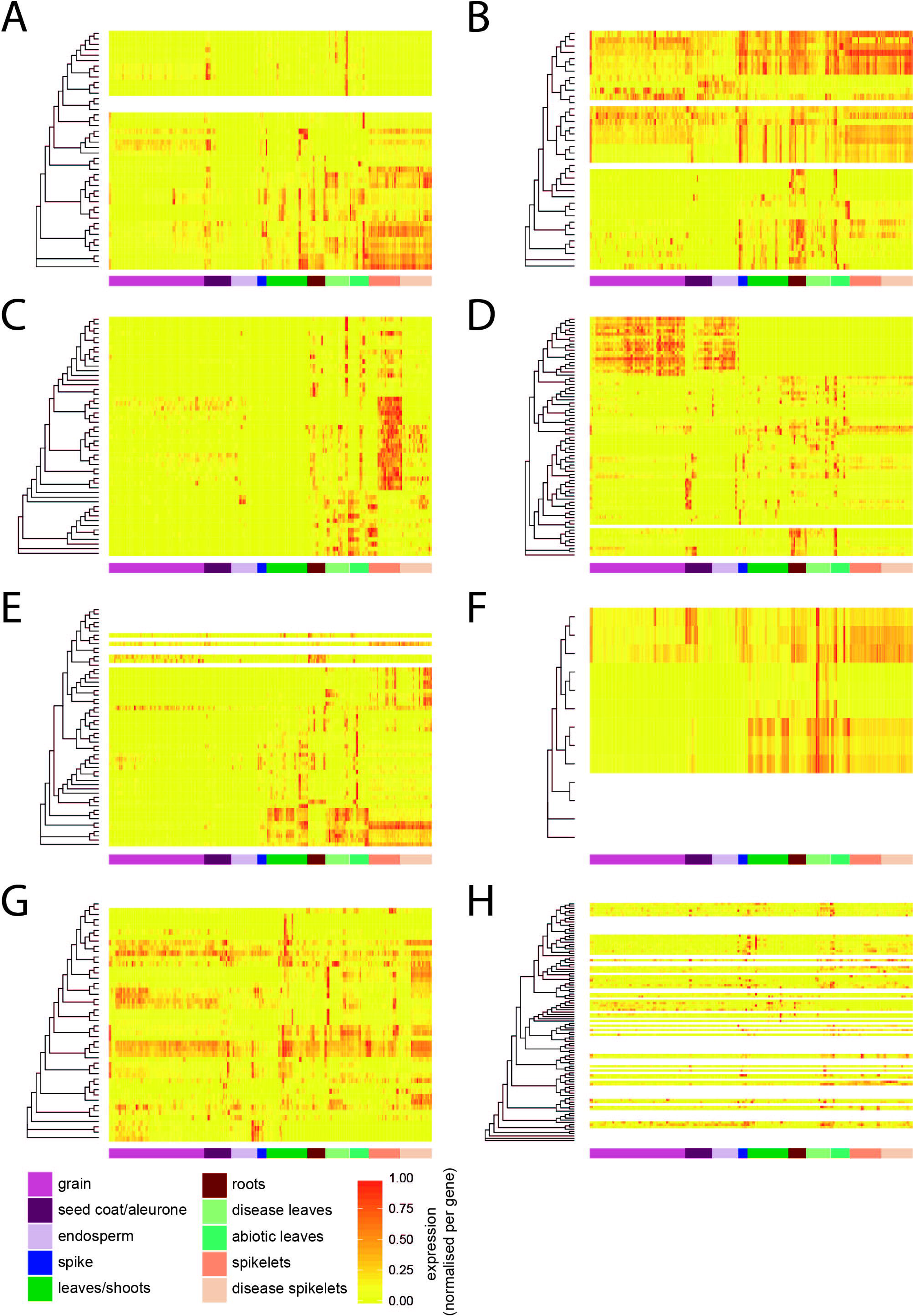
Relationship between phylogenetic position and NAC gene expression across 308 RNA-seq samples from diverse tissues, developmental stages and stress conditions. The origin of each sample is indicated by the colored bar under each heatmap. Each panel (A-H) represents NAC genes belonging to that group according to the classification in Figure 2. Dendrograms indicate the maximum likelihood phylogeny of genes within each group. Genes which did not meet the minimum expression criteria (>0.5 tpm in at least three samples) do not have expression data represented (white rows). All remaining expression data (tpm) was normalized per gene to range from 0 to 1. An extended version of the figure with the full phylogenetic trees is available as Figure S3.

To explore the patterns of NAC TF expression in a global context we carried out co-expression analysis using Weighted Gene Correlation Network Analysis (WGCNA) across all gene families using the 308 RNA-seq samples. We could assign 61,325 genes (out of 91,403) to 37 co-expression modules (clusters) which ranged in size from 46 to 11,082 genes with a mean size of 1,546 genes (Figure 5A) (Table S7). In total 3,446 TFs (out of 5,776) were assigned to modules and these made up on average 5.9 % of genes within each module (Figure 5B). In total 259 NACs (out of 453) were assigned to 23 of the 37 modules (Figure 5C). NAC TFs were over-represented (χ^2^ p<0.05) within modules 1, 6, 20, 29 and 34, respectively as 11%, 12 %, 17 %, 31 % and 21 % of all TFs in those modules were NACs compared to an average across all modules of 8 %. We carried out GO term enrichment on all genes within these modules and found these modules are enriched for phosphorylation (module 1), exocytosis and cell wall organization (module 6), protein export from nucleus and response to water (module 20), photosynthesis (module 29) and regulation of photoperiodism and flowering (module 34) (Table S8). In general NACs within co-expressed modules were from several phylogenetic groups (Figure 5D). However certain modules e.g. 17, 20, 26 and 29 contained genes from only one group (b, d, c and a, respectively). These modules were enriched for GO terms related to response to heat and abiotic stress (module 17), protein export from nucleus and response to water (module 20), protein phosphorylation and system development (module 26) and photosynthesis (module 29).

**Figure 5.**
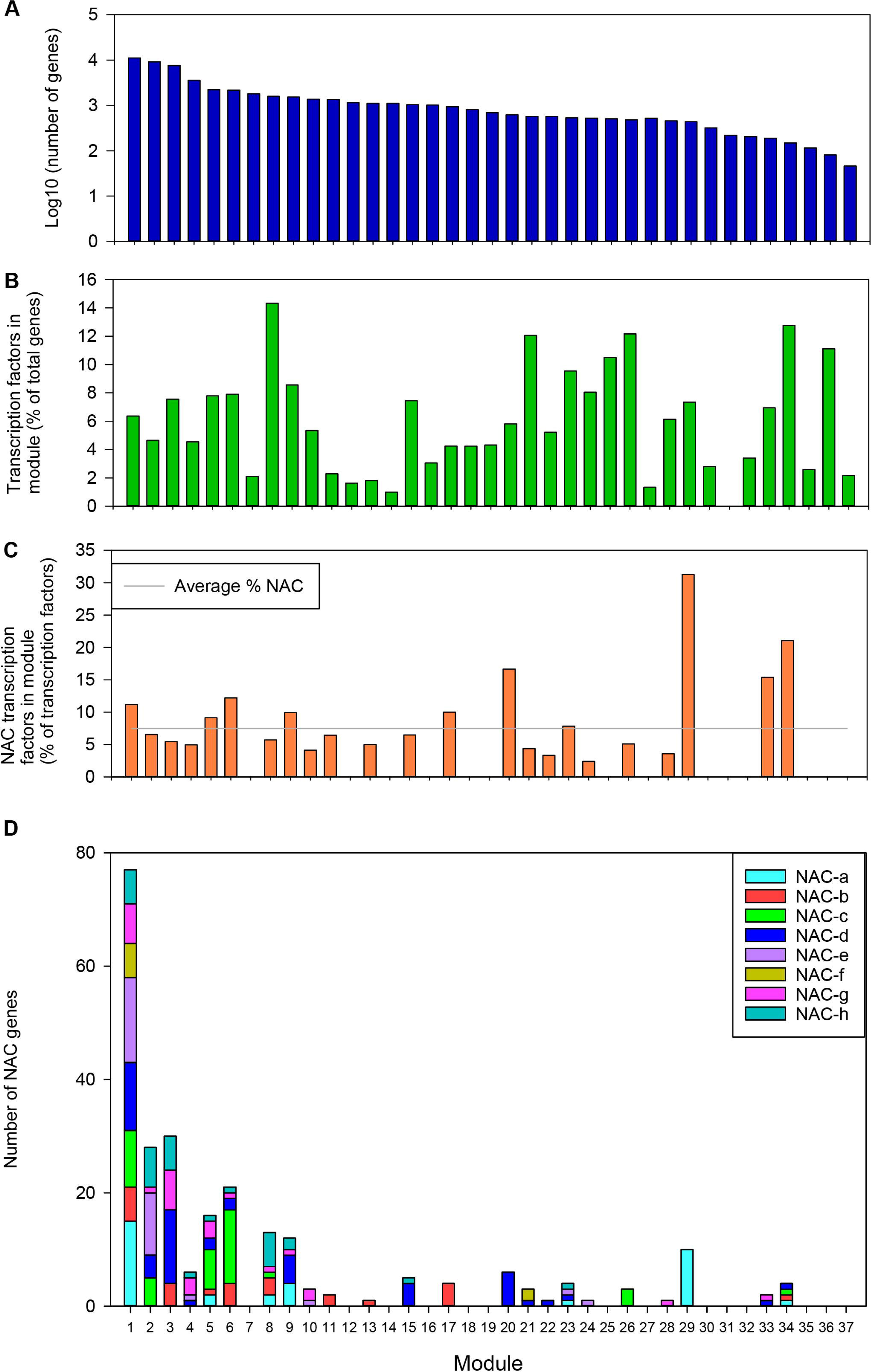
Distribution of genes and TFs across modules. A) Number of genes (log10), B) Percentage of genes which are TFs, C) Percentage of TFs which are NAC TFs, D) Number of NACs from each phylogenetic group.

This indicates that some phylogenetically related NACs share similar expression profiles and may be involved in regulating similar biological processes. Interestingly module 20 and 29 were both enriched in NACs compared to other TFs and specifically in NACs from groups d and a, respectively. This indicates that NACs may play a relatively major role in the regulation of these processes given their over-representation compared to other TFs in these co-expressed modules.

## DISCUSSION

The availability of a more complete genome sequence for wheat has allowed the comprehensive analysis of wheat TF families. We identified 5,776 TF genes which is 1.5-3 fold higher than has previously been reported for wheat (3,820 in wDBTF (Romeuf et al. 2010), 2,407 in WheatTFDB (Chen et al. 2015) and 1,940 in PlantTFDB (Jin et al. 2014)). We found that overall wheat has 5.1 times more TFs than in other diploid Triticeae species. Compared to rice, wheat has 3.1 times more TFs as would be expected for a hexaploid species. The incomplete nature of the genomes of other monocots may explain the higher than expected ratio (3:1) of wheat TFs to monocots other than rice. Each family is present in wheat in similar proportions to those found in other monocots. The NAC family is one of the largest TF families and has been characterized previously in other species (Ooka et al. 2003; Christiansen et al. 2011; Nuruzzaman et al. 2010; Peng et al. 2015; Saidi et al. 2017; Le et al. 2011). This is, however, the first study to identify the NAC genes in hexaploid wheat and characterize their global expression patterns. We found that NAC TFs were located across all chromosomes, but were most frequently found on chromosome group 2 (on average 39 NACs per homoeolog) with relatively few NACs on group 1 (on average 7 NACs per homoeolog). The uneven distribution of NACs across chromosomes has also been observed in rice (Nuruzzaman et al. 2010) and maize (Peng et al. 2015).

We found that wheat NAC TFs belong to eight main phylogenetic groups, similar to Arabidopsis, rice and barley. Wheat has a reduced f group with only 13 genes compared to the 10 genes found in rice, but not compared to barley suggesting that group f NACs were reduced in number in the ancestral Triticeae. This family specific reduction requires further investigation to determine its biological relevance.

The DNA and protein-binding NAC domain of NAC TFs has been studied over the past two decades (Xie et al. 2000; Welner et al. 2012; Ernst et al. 2004), however the function of the C-terminal domain remains poorly understood. We detected previously identified CTD motifs in wheat, rice and barley NAC TFs and also identified three novel CTD motifs. These motifs were in general restricted to one or two NAC groups. The presence of these motifs was typically conserved within closely related clades of rice, barley and wheat orthologues. This is expected given the high overall sequence similarity between orthologs in these species. However, the conservation of CTD motifs extends beyond the immediate orthologs in these species. For instance, motifs iii and xiii in group c are conserved across several discrete clades which contain rice, barley and wheat members. This evolutionary conservation inside otherwise non-conserved regions indicates that CTD motifs may have important biological functions. The *de novo* identification of CTD motifs which match those identified in studies of other plant species also highlights the conservation of motifs within angiosperms and indeed the plant kingdom as a whole (Pereira-Santana et al. 2015; Shen et al. 2009; Ooka et al. 2003). These motifs are thus good candidates for further investigation into the role of the NAC CTD and the specific function of these motifs.

In this study we also combined global gene expression data from 308 RNA-seq samples with TF annotations. We found that within the phylogenetic groups a-h there are variations in expression patterns although there are clades of genes which have extremely similar patterns. These genes with conserved expression patterns in particular tissues may represent good candidates to explore for functional roles in those tissues. In rice, for example, the use of co-expression as a guide to putative function has been successful in identifying several transcription factors regulating grain filling (Xu et al. 2016; Fu and Xue 2010) suggesting this method might also prove useful in wheat. Sequenced mutant populations (Krasileva et al. 2017) and gene editing methods (Wang et al. 2014; Liang et al. 2017; Zhang et al. 2016) provide a direct route for hypothesis testing.

We produced co-expression modules which can be used to inform a range of further studies. Focusing on wheat NAC TFs we found several examples where GO term enrichment of co-expressed genes supports known TF function. For example, *TaNAC-S* was found to be co-expressed with genes related to photosynthesis (module 2) according to GO term enrichment. It has previously been shown that *TaNAC-S* over-expression delays senescence and increases the expression of Rubisco which a central enzyme for carbon fixation in photosynthesis (Zhao et al. 2015). *TaNAM1* and *TaNAM2* were found in module 9 which is enriched for protein ubiquitination related genes. *TaNAM* genes are known to increase protein content in the grain by increasing the remobilization of nitrogen from vegetative tissues (Waters et al. 2009). The ubiquitin pathway has previously been linked to senescence (Vierstra 2003) and several e3 ubiquitin ligases are downregulated in *TaNAM1* and *TaNAM2* mutants (Pearce et al. 2014) indicating these genes may act through the ubiquitin pathway to bring about protein degradation for remobilization during senescence. Several NAC TFs including *TaNAC2* and *TaNAC4* have been reported to be responsive to both abiotic and biotic stresses (Xia et al. 2010a; Mao et al. 2012; He et al. 2015) and their co-expression with genes involved in protein phosphorylation (module 1) may provide a putative mechanism as to how they regulate responses to multiple stresses. These examples indicate that our coexpression modules categorize known genes with appropriate GO terms. GO term enrichment may also be predictive of the functions of novel genes (Eisen et al. 1998). For example in *Arabidopsis thaliana* a zinc finger transcription factor (*AtZFP2*) was predicted to regulate abscission due to its expression within a group of genes which had GO terms associated with cell wall modifying proteins, extracellular regulators, and transcription factors. *AtAFP2* was subsequently demonstrated to regulate abscission in overexpression lines (Cai and Lashbrook 2008).

Previously characterized wheat NAC TFs were only identified in five co-expression modules out of the total 23 modules in which NAC TFs were expressed. This indicates that NAC TFs may still play unrecognized roles in wheat. This study provides the framework for further investigations of NAC TF function in this important crop species.

## ACKNOWLEDGEMENTS

We thank Alejandro Pereira for sharing sequence data used for MEME analysis in his previous publication (Pereira-Santana et al. 2015). We thank Per L. Gregerson for providing a list of barley NACs (Christiansen et al. 2011) updated with MLOC nomenclature. This work was funded by the UK Biotechnology and Biological Sciences Research Council (BBSRC) Grants BB/P013511/1 and BB/P016855/1, BBSRC Future Leader Fellowship BB/M014045/1 to P. Borrill, and the John Innes Foundation.

